# Effects of the Automatic Self-Transcending Meditation on cognition and mental states in the EEG, skin conductance and behavioral performance: a pilot study

**DOI:** 10.1101/2022.10.11.511756

**Authors:** Lucas Galdino, Gabriella Medeiros Silva, Thiago A. S. Bonifácio, Natanael Antonio dos Santos, David Orme-Johnson

## Abstract

The main purpose of this pilot study was to investigate the immediate effects of Automatic Self-Transcending (AST) meditation on the cognitive function, EEG activity and autonomic arousal (Study 1) and characterize the frontal EEG synchrony during resting state, cognitive activity and AST mental states in traditional and wireless EEG’s (Study 2). We report the results of three healthy AST meditation volunteers in this case-report study (Case 1 - age = 26 years, meditative practice time = 2 months; Case 2 - age = 39 years, meditative practice time = 6 years; Case 3 - age = 59 years, meditative practice time = 40 years). In study 1, the volunteers performed a protocol with simultaneous recording of EEG and skin conductance while performing the Stroop test (T0), followed by 20 minutes of AST meditation and immediately the same protocol performed at T0 (T1). We analyzed P300 amplitude and latency, as well as test behavioral response and skin conductance activity before and immediately after a single session of AST. In study 2, the same volunteers performed three tasks with eyes closed on two EEG equipment (traditional and wireless): resting state, cognitive activity and one session of AST, each for 20 minutes. We measured the frontal interhemispheric coherence of alpha1 (8-10 Hz) and beta (13-30 Hz) for each condition and EEG type. Our main findings show that there is an immediate beneficial effect after AST meditation at the level of the same individual with different patterns of P300 and skin conductance activity and that AST meditation is marked by an overall increase in the frontal coherence of alpha1 and beta bands, when compared to other mental states. We conclude that 1) there is an immediate effect on cognition and executive control after AST meditation, 2) the frontal interhemispheric coherence of alpha1 and beta bands are increased during AST, and 3) wireless EEG exhibits the same characteristics observed in traditional EEG and therefore can be used to describe cortical dynamics during AST.

## 1. Introduction

Automatic self-transcending (AST) is an effortless category of meditation characterized by a state of transcendence mediated by minimal cognitive control (Travis, 2020) that creates a transitional state of consciousness that is not mediated by voluntary focused attentional processes (Travis & Shear, 2010). This meditation technique can be also defined as part of a null-directed methods (NDM) category which includes Transcendental Meditation (TM), Zen satori and some Yoga methods (Nash et al, 2013). This study was on TM, the most studied with robust results on its electrophysiological biomarkers and lifetimes benefits, such as cognitive intelligence, autonomic stability and emotional reasoning (Orme-Johnson, 2021). The subjective experience during the AST meditation is marked when thoughts are “Stilled” and “Pure Awareness” is gained as a result of the absence of sensory and mental contents (Travis, 2014). This state is characterized by a dissolution of the sense of the localized self and leads the meditator to a state in which he/she does not differentiate the self and the exterior world during the practice (Deolindo et al., 2020).

TM has been taught worldwide in a standardized format by certified teachers since the early 1960’s (Maharishi Mahesh Yogi, 1963). One learns a suitable mantra and how to use it effortlessly. During the practice, the mind is automatically drawn inward via the mantra settling to subtler levels of thoughts. Thus, the mind automatically transcends or goes beyond the process of the technique itself (the mantra) to a quiet space of no-mantra and no thoughts.

The electroencephalography (EEG) activity during TM includes an increased alpha 1 (8-10 Hz) power (Travis, 2001a) and coherence (Travis, 2001b; Hebert et al., 2005; Travis et al., 2010) over the frontal cortex is well documented and shows that during the TM practice the local oscillation of the alpha1 band may reflects an enlargement of interhemispheric phasic connection (i.e., Magnitude Squared Coherence). A review of EEG studies on meditation found more than a dozen studies reporting increased frontal alpha1 power and coherence during TM (Cahn & Polich, 2006). Parallel to EEG results, a functional magnetic resonance imaging study shows that the blood oxygenation level dependent (BOLD) is increased in frontal areas, followed by a decrease in the pons and brainstem BOLD, corresponding to an enlivenment of executive process that are associated with enhanced awareness and decreased motor activity, respectively (Mahone et al., 2018).

More recent studies show that these changes observed in the frontal cortex come from important subregions that mediate cognitive processing and memory, for example. Yamamoto et al., (2006) showed that medial prefrontal cortex (mPFC) and anterior cingulate cortex (ACC) plays an important role in the findings with EEG. The mPFC has a regulatory role in numerous cognitive functions, including attention, inhibitory control, habit formation and working, spatial or long-term memory (Euston et al., 2012). The ACC has roles in many cognitive processes, integrating and modulating the emotional reactions to experience as in empathy, impulse control, emotion, decision-making, and social behavior (Apps et al., 2016).

However, most studies have been conducted with traditional EEG (T-EEG) devices in settings that do not directly reflect the daily practice of the meditators in their natural environments. Rather, most of our knowledge of meditation is from studies in artificial controlled laboratory environments such as universities and private meditation centers. In this context, Geers et al., (2005) and Price et al., (2008) postulated the motivational effect in studies involving brain measurements performed in a controlled environment may act like a confounding variable and possibly enhance the observed results. In an ideal setting to describe how interhemispheric connectivity occurs during the daily practice of AST meditation, it is necessary to conduct an investigation using a device that is wirelessly portable and user friendly. The Emotiv X wireless EEG (W-EEG) is a commercially-made low-cost electroencephalography (EEG) device that has become increasingly available over the last decade, thanks to the ease in acquiring EEG signals outside the traditional laboratory setting (Ekanayake et al., 2010; Stysenko et al., 2011; Badcock et al., 2013; Boutani & Ohsuga, 2013). However, this device has not yet been validated for such purposes, and also similar results with portable EEG’s were never described yet as well as the investigation of the immediate effects of just one AST meditation session on cognition and impulse control.

To answer these questions, we design two case-report pilot studies which aim to measure the immediate effects of AST meditation on the cognitive function and autonomic arousal (Study 1) and characterize the frontal EEG synchrony during resting state, cognitive activity and AST mental states in traditional and wireless EEG’s (Study 2).

## 2. Material and methods

A randomized, quasi-experimental, and single-blinded trial with repeated measures was conducted during two consecutive days. We report the results of three healthy AST meditation volunteers in this pilot study (Case 1 - age = 26 years, meditative practice time = 2 months; Case 2 - age = 39 years, meditative practice time = 6 years; Case 3 - age = 59 years, meditative practice time = 40 years). All procedures performed were approved by the Research Ethics Committee of the Federal University of Paraíba, Brazil (Registration number: 69737817.4.0000.5188) and followed the ethical principles of the Declaration of Helsinki.

### 2.1. Electroencephalography (EEG)

#### 2.1.1. Traditional EEG (T-EEG)

We use the actiCHamp EEG system (Brain Products, Herrsching, Germany) to access the cortical activity. A sampling rate of 1000 points per second was used with 32 active electrodes (Fp1, Fz, F3, F7, FT9, FC5, FC1, C3, Cz, T7, TP9, CP5, CP1, Pz, P3, P7, O1, Oz, O2, P4, P8, TP10, CP6, CP2, C4, T8, FT10, FC6, FC2, F4, F8, and Fp2) distributed according to the 10/5 system placed in an air permeable cap adjustable to each participant’s head (Easy-cap, Herrsching, Germany). The reference and ground electrodes were placed at Pz and Cz levels, respectively. In addition, a saline gel solution (Easy-cap, Herrsching, Germany) was used to promote the physical contact between the electrode and the scalp and sustain the impedance below 10 kΩ during the experiment.

#### 2.1.2. Wireless EEG (W-EEG)

The Emotiv Insight X is a wireless portable EEG device with 14 active electrodes (AF3, F7, F3, FC, T7, P7, O1, O2, P8, T8, FC6, F4, F8, and AF4) displaced over the temporal, parietal, frontal and occipital lobes that follows the 10/20 international positioning system. We used the default specification of the device: sampling rate of 128 points per second, and no analog filter. Prior to the recordings, the electrodes were embedded in a NaCl solution to promote contact with the scalp and establish a low impedance during the entire experiment.

### 2.2. Auto-Self Transcending meditation

All volunteers performed at least one AST session in the two proposed studies. The session was held in a controlled laboratory experimental environment that could be performed with T-EEG or W-EEG for 20 minutes following the meditation protocol previously used by the volunteers. Meditation sessions for both experiments were conducted using an [AST guided meditation mobile app](https://apps.apple.com/us/app/v%C4%ABrya-guided/id1562147704) that timed the start and end of the meditation with audible instructions designed specifically for AST meditation. The AST meditation used in this study was the standard Transcendental Meditation technique (Elder et al., 2014).

### 2.3. Study 1 - Immediate effects of AST meditation on the cognitive function and autonomic arousal

In study 1, we used the simultaneous recording of the EEG with Skin Conductance (SC) during the Stroop test immediately before and after an AST session. Prior to the beginning of the experiment, the screening was performed to assess whether the subjects met the study’s inclusion criteria and then the informed consent was signed. After the screening, the subjects performed the three steps proposed in the study (Figure 1AB): 1) T0 period where the subjects performed the Stroop test with simultaneous recording with T-EEG and SC for approximately 16 minutes; 2) A single session of AST for 20 minutes with closed eyes in a dark room with controlled temperature; 3) T1 period, the same task proposed in T0 to measure the acute and immediate effects of AST. The procedures for analyzing EEG, SC and behavioral data from the Stroop test are illustrated in Figure 1C

**Figure 1.**
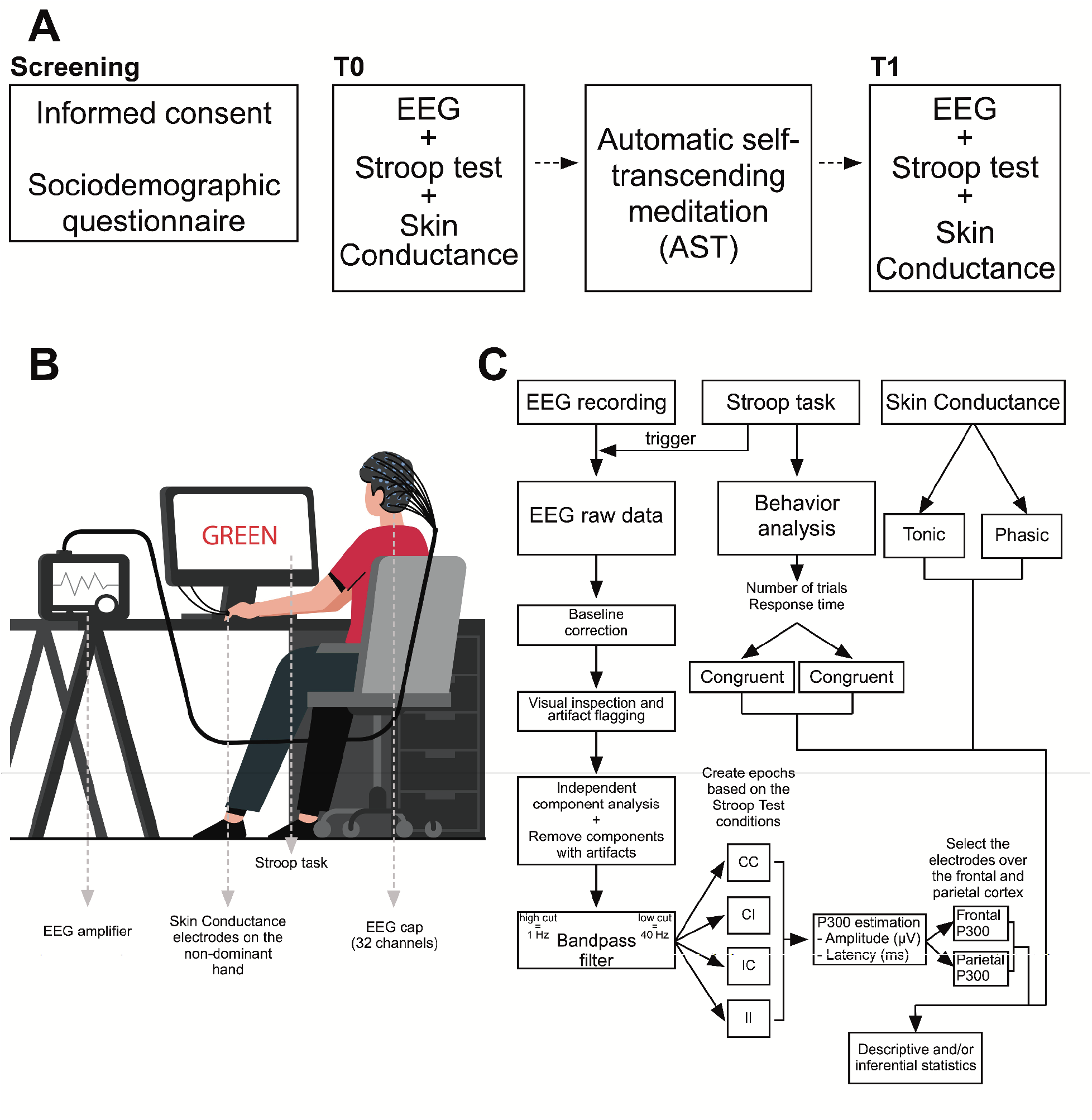
Flow of the experimental design proposed for the study 1. A) General experimental setup showing the screening section, pre (T0) and post (T1) measurements taken immediately before and after a single 20-minute AST session. B) Illustration of the arrangement of each device (EEG and SC) during the Stroop test at T1 and T2. C) Diagram showing the EEG, SC and behavioral processing steps of the Stroop test performed offline for each study volunteer.

#### 2.3.1. EEG analysis

EEG datasets were processed using EEGLAB (Delorme & Makeig, 2004) and custom-made scripts with the following steps (Figure 1C): 1) baseline correction, 2) visual inspection and artifact removal, 3) Independent Component Analysis (ICA) followed by inspection of the 32 components through the ICLabel, an algorithm that shows the correlations of each component with artifacts such as eye movement, muscle activity, and electrical noise (Pion-Tonachini et al., 2019), 4) removal of ICA components with artifacts, 5) butterworth bandpass filter with order 4 between 1 and 40 Hz, 6) data epoching using the photo sensor trigger with 200 ms of prestimulus interval and 600 ms of event-related potential activity, 7) separation of the epochs according to the conditions of the Stroop test, 8) identification of the highest peak between 250 and 350 ms and its respective latency, 9) selection of electrodes over the frontal (Fp1, Fz, F3, F7, F4, F8, and Fp2) and parietal cortex (Pz, P3, P7, P4, P8), 10) statistical analysis of P300 amplitude and latency in the frontal and parietal regions p for each condition of the Stroop test.

#### 2.3.2. Stroop Task

A computer version of the Stroop test was applied using the Psychopy software (v. 3.0). The stimuli were three color names presented in upper-case letters as red, blue and green in two conditions: congruent and incongruent. In the congruent condition, meaning of the word and its color matched, and in the incongruent condition, the color of the font was different than the word meaning (Stroop, 1935; Scarpina & Tagini, 2017). All stimuli were presented with 3.0 × 2.5 cm letters in the center of a 13 inch computer screen over a gray background at 60 cm distance. We applied 100 congruent and 100 incongruent trials in randomized order. Participants’ tasks were to press one of the three buttons on the computer mouse as follows: left button for red color, middle button for green, right button for blue with their index finger, middle finger and ring finger, respectively with the dominant hand (right hand). Each stimulus appeared on the screen until the participant responded with an interstimulus interval of 1500 ms. The trigger for the EEG epoching was synchronized using a photosensor (Brain Products, Herrsching, Germany) plugged on the screen which sent a binary signal on an auxiliary electrode on the EEG with the same sampling rate specifications and time-synchronized. Behavioral analysis was performed offline and the number of trials and response time for the congruent and incongruent conditions were measured.

#### 2.3.3. Skin Conductance (SC)

The skin conductance was measured from the proximal phalanges of the index and the middle fingers of the non-dominant hand at a frequency of 20Hz using a Neulog NUL-217 device. The participants were asked to keep their hands relaxed on their thighs to minimize the influence of movement on the measured signals and the environment temperature was maintained at 64.4–68?F to minimize the effect of temperature. The signals were processed digitally and offline and decomposed into tonic and phasic signals to better describe the evoked skin conductance during the stroop task (Lim et al., 1997).

### 2.4. Study 2 - Characterization of frontal EEG synchrony during AST in traditional and wireless EEG

EEG recording was done with both devices (T-EEG and W-EEG) one day after Study 1. In Study 2, the participants’ task was to perform three consecutive recordings in each type of EEG with the following experimental conditions (Figure 2A): resting state (RS), performing one mental calculation task that consisted of counting from 1000 to 1 in steps of 4 (Cognitive Activity or CA) and performing an AST session (AST). The device type sequence was previously mixed for each subject, while the order of the experimental conditions was chosen by each subject, configuring a blind study for the experimenter who did not know which sequence was performed in each experimental session. All experimental conditions in both devices were carried out in the same sound proof room with controlled temperature and humidity.

**Figure 2.**
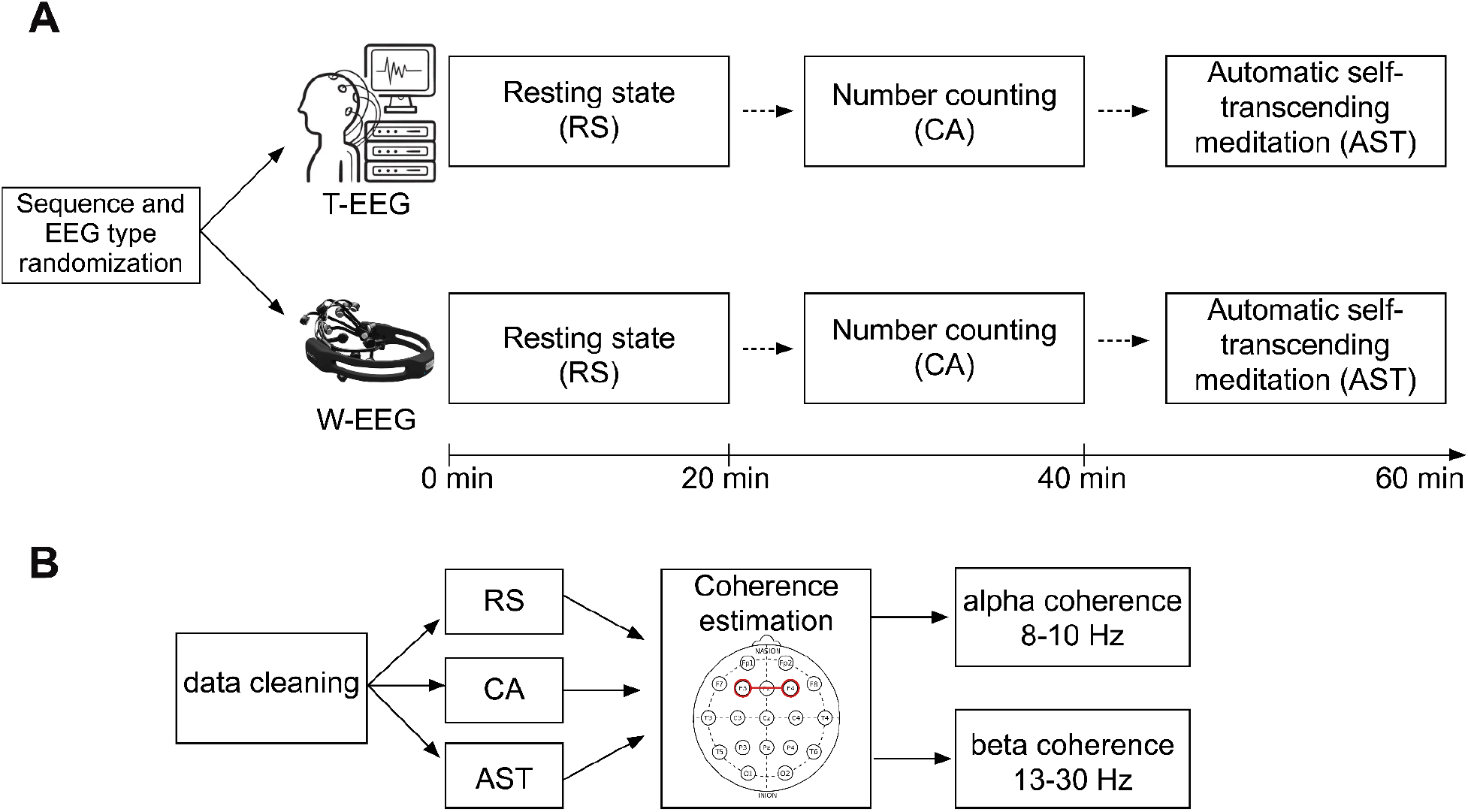
Experimental design of the Study 2. A) The EEG type and the single-blinded experimental conditions performed during the experiment. B) EEG analysis and coherence estimation between F3-F4 electrodes and its spectrum segmentation into alpha and beta coherences.

#### 2.4.1. EEG analysis

The blind analysis (Figure 2B) of each experimental condition (RS, CA or MRM) was performed for each type of EEG (traditional or wireless) following the same steps performed in study 1 until the fifth point. Thus, from data cleaning using ICA and visual artifact removal, the coherence measure by phase connectivity (Equation 1) was applied between the pair of front electrodes common to both the W-EEG and the T-EEG (F3-F4) during the 20 minutes of each condition. Coherence estimates the phase coupling between x and y signals at a frequency (f) by measuring the squared magnitude of the complex cross power spectral density (Pxy) divided by the auto-spectral densities (Pxx) and (Pyy) (Bowyer, 2016). We used the following specifications on the native function *mscohere* implemented in the MATLAB: window size of 4 * sampling rate with 50% overlapping and NFFT of 2048 points. Spectrum coherence was then averaged in the low-alpha (or alpha1) and beta frequency bands, which consist of the 8-10 Hz and 13-30 Hz frequency ranges, respectively.

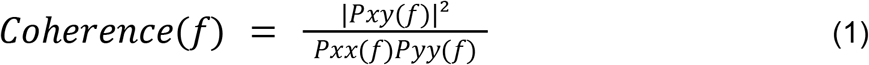

## 3. Results

The results described below are separated for each case proposed in the study using descriptive statistics for the variables of the behavioral analysis of the Stroop test, amplitude and latency of the P300 component in different cortical regions and SC. Yet, paired inferential analysis was performed using the Student’s t-test corrected for multiple comparisons in the electrophysiological study during the Stroop task, as the data were segmented into epochs and thus it becomes possible to obtain a sufficient sample size to perform this test. The evoked-response plots for experiment 1 is shown in Supplementary Material (S1).

### 3.1. Case 1

#### 3.1.1. Study 1

In the behavioral analysis of the Stroop test, the first case showed a reduction in the response time (Figure 3A) for the conditions CC (T0 mean = 0.75; T1 mean = 0.61), CI (T0 mean = 0.53; T1 mean = 0.49), IC (T0 mean = 0.81; T1 mean = 0.72), II (T0 mean = 0.74; T1 mean = 0.60). The electrophysiological properties of the P300 component showed statistic differences for the incorrect trials in the congruent and incongruent trials of the Stroop task. In detail, the amplitude modulation of the CI trials after a single session of AST was observed over the frontal electrodes (Figure 3B) between T0 (mean = 3.74; standard error of the mean (sem) = 0.58) and T1 (mean = 9.30; sem = 23.4) with confirmed statistical difference observed in the two-tailed student’s T test corrected for multiple comparisons (t = 2.06; *p* = 0.04), and also for the parietal (Figure 3C) response T0 (mean = 2.13; sem = 9.99) and T1 (mean = 15.2; sem = 3.32) (t = -3.81; *p* < 0.01). We also observed that the P300 latency was significantly decreased over the frontal electrodes (Figure 3B) for II trials between T0 (mean = 310.4; sem = 2.27) and T1 (mean = 301.03; sem = 30.75) (t = 2.77; *p* < 0.01), and the parietal electrodes (Figure 3C) for IC trials T0 (mean = 321.31; sem = 26.22) and T1 (mean = 310.86; sem = 3.54) (t = 2.43; *p* = 0.01).

**Figure 3.**
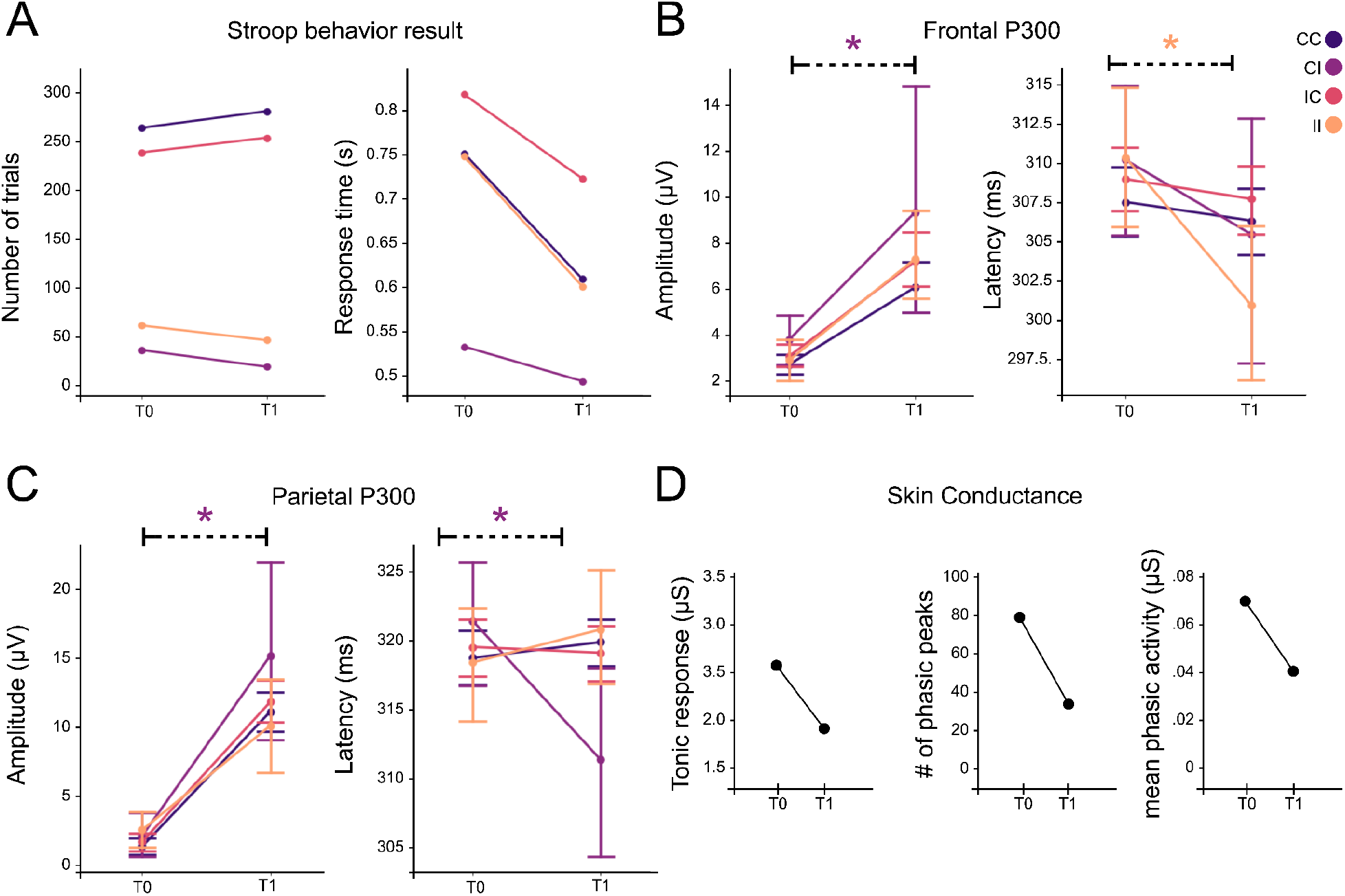
Case 1 results for experiment 1. The colors represent the Stroop task conditions (CC - Congruent Correct, CI - Congruent Incorrect, IC - Incongruent Correct, and II - Incongruent Incorrect). * show the statistical significance (two-tailed student’s T test corrected for multiple comparisons) < 0.05. The error bars represent the standard error of the mean for each condition.

The SC analysis between T0 and T1 showed that this case exhibits a global decrease in the skin conductance activity (Figure 3D). We observed a decrease in the tonic response (T0 mean = 2.67; T1 mean = 2.03), number of phasic peaks (T0 peaks = 81; T1 peaks = 40) and mean phasic activity (T0 mean = 0.07; T1 mean = 0.04).

#### 3.1.2. Study 2

In the second study we compared different mental stages between two EEG devices. The case 1 results showed that regardless of the EEG type (traditional or wireless), the frontal coherence is increased during the AST session. Therefore, this volunteer showed the following coherences for both alpha in T-EEG: RS = 0.17, CA = 0.39, and AST = 0.69, following the same pattern in W-EEG: RS = 0.22, CA = 0.21, and AST = 0.56. Beta coherences showed the same pattern of increased frontal coherence during AST for both T-EEG (RS = 0.22, CA = 0.40, and AST = 0.70) and W-EEG (RS = 0.08, CA = 0.14, and AST = 0.39).

### 3.2. Case 2

#### 3.2.1. Study 1

The stroop task behavior analysis for the case 2 (Figure 5A) showed that in T1 the response time was lower for the conditions CC (T0 mean = 0.90; T1 mean = 0.68), IC (T0 mean = 0.82; T1 mean = 0.71), II (T0 mean = 0.75; T1 mean = 0.71) and higher only for CI trials (T0 mean = = 0.68; T1 mean = 1.28). The frontal P300 analysis showed statistic difference only for II trials with a increase in the amplitude (T0 - mean = -8.82; sem = 45.62 and T1 - mean = 3.93; sem = 1.04; t = -1.98; *p* = 0.05) and a decrease in the latency after AST (T0 - mean = -8.82; sem = 45.62 and T1 - mean = 3.93; sem = 1.04; t = -1.98; *p* = 0.05). The parietal response showed a decrease in the P300 amplitude in T1 (T0 - mean = 11.16; sem = 35.33 and T1 - mean = 7.52; sem = 0.43; t = 3.16; *p* < 0.01) and no significant difference for the parietal P300 latency. The skin conductance response showed that the case 2 exhibit a decrease in the skin conductance activity (Figure 3D) in the tonic response (T0 mean = 2.04; T1 mean = 1.83), number of phasic peaks (T0 peaks = 17; T1 peaks = 11), while an increase in mean phasic activity was observed (T0 mean = 0.03; T1 mean = 0.06).

#### 3.2.2. Study 2

The analysis of the second study for case 2 showed results similar to those observed for case 1. On the T-EEG the frontal coherence is increased in the AST condition in both alpha (RS = 0.35, CA = 0.64, and AST = 0.80), and beta (RS = 0.46, CA = 0.60, and AST = 0.67). In addition, the T-EEG result shows the same pattern for alpha (RS = 0.08, CA = 0.13, and AST = 0.54) and beta (RS = 0.14, CA = 0.17, and AST = 0.46) (Figure 6).

### 3.3. Case 3

#### 3.3.1. Study 1

The behavior analysis during the Stroop task showed the same pattern observed in the case 1, a global decrease of response time for all conditions in T1 (Figure 7A): CC (T0 mean = 0.73; T1 mean = 0.68), CI (T0 mean = 0.76; T1 mean = 0.66), IC (T0 mean = 0.81; T1 mean = 0.75), II (T0 mean = 0.84; T1 mean = 0.68). On the other hand, a different electrophysiological signature was observed for the P300 component in the frontal P300 (Figure 7B). We found a difference in amplitude in the CI trials between T0 (mean = 3.54; sem = 7.96) and T1 (mean = 39.95; sem = 59.36) (t = -2.15; *p* = 0.03). No differences were found between the frontal latency and the P300 activity over the parietal cortex.

The SC activity showed an overall pattern that was not expected and did not satisfy our hypothesis (Figure 7D). We found that the tonic response (T0 mean = 2.30; T1 mean = 3.03), number of phasic peaks (T0 peaks = 37; T1 peaks = 54), and the mean phasic activity (T0 mean = 0,041; T1 mean = 0.048) is higher after the AST session, showing that for this case the meditation practice elevated the skin conductance during a cognitive task such as the Stroop Task in a different manner.

#### 3.3.2. Study 2

As expected, the study 2 results showed results similar to those observed for case 1 and case 2 (Figure 8). In detail, the T-EEG frontal coherence is increased in the AST condition in both alpha (RS = 0.40, CA = 0.42, and AST = 0.66), and beta (RS = 0.45, CA = 0.32, and AST = 0.56) as well as in the W-EEG for alpha (RS = 0.31, CA = 0.47, and AST = 0.55) and beta (RS = 0.29, CA = 0.44, and AST = 0.53) bands (Figure 8).

## 4. Discussion

### 4.1. Study 1 - Immediate effects of AST meditation on the cognitive function and autonomic arousal

We investigated the immediate effects of AST meditation on the cognitive behavioral performance of the Stroop test associated with simultaneous recording of EEG and SC. In the EEG, we analyzed the amplitude and latency of the P300 component in the parietal and frontal electrodes for each condition of the Stroop test, while for the SC we analyzed the tonic response, number of phase peaks and mean phasic activity during the entire experimental task before (T0) and after (T1) a single session of AST meditation. Our results confirm the initial hypothesis of an immediate improvement in the behavioral result of the Stroop test; however, we found a different and unique pattern of EEG event-related activity and skin conductance (tonic and phasic) for each case reported in the study. Thus, the results indicate that a global decrease of time response for all conditions during the task was observed, except for the CC trials of the case 2. The electrophysiological response for case 1 showed that immediately after the AST meditation the amplitude and latency of the P300 only for CI and II conditions in the parietal and frontal region was modulated. On the other hand, the second case showed changes in amplitude and latency in the frontal P300 in the II condition, whereas in the parietal region a significant decrease in the amplitude of the P300 was observed only for the CC condition. Interestingly, case 3 showed only an increase in frontal P300 amplitude for the CI condition.

The P300 in the parietal cortex is related to the attentional component to elucidate a specific type of stimulus and, therefore, it is related to the attention process mediated by an increase in sensory integration (Kropotov, 2009). Other authors define it as a result of the synchronization of neurons that control selective attention (Polich, 2007; Pontifex, Hillman, & Polich, 2009) via direct efferent pathways from the locus coeruleus, a key region to attentional, arousal and emotional processes which sends projections to the parietal and frontal cortex. In addition to evidence related to attentional processes, the parietal P300 is also related to cognitive processes, such as learning (Bouret, 2004), memory (Uematsu, 2015) and adaptation/behavioral flexibility (Aston-Jones, 1999). The role of the frontal P300 is less studied, but strong evidences indicate that this component is associated with cognitive and inhibitory control (Kim, 2001; Kopp et al., 2006; Gajewski, 2017) and stimulus-driven frontal attention (Polich, 2007) which may be linked to impulsive behaviors and decision making (Harmon-Jones, 1997; Barceló, 2003), being an important complementary measure during the Stroop test and also to measure the immediate effects of AST.

Still about the role of autonomic arousal during the Stroop task, the SC is an important measure of how we evoke stimulus-driven signatures of stress or emotional states through the continuous variation in the electrical characteristics of the skin (Montagu & Coles, 1966; Critchley et al., 2000; GerŠak et al., 2012). To do so, we segmented the raw signal into tonic signal — related to lower oscillations —, and phasic signals — related to arousal evoked by an external stimulus, which in our case are the four different conditions of the Stroop test. The results indicate that each case reported in the study has a different SC response, which may be related to the electroencephalographic measurement of P300. For case 1, an overall decrease was observed in all skin conductance variables (Figure 3D), while case 2 was marked by a decrease in tonic activity and number of phasic peaks and an increase in mean phasic activity (Figure 5D). However, the case with the greatest number of years of meditation obtained a different result from our initial hypothesis: an increase in SC activity for all measures (Figure 7D).

An important point that needs to be addressed in the replicability of the Stroop Test in the same sample. The test retest reliability is a measure of reliability obtained by administering the same test twice over a period which aims to describe the stability of the cognitive assessments over time (Cronbach, 1971). It is well described that certain aspects of the Stroop Test is affected by a simple retest, but the majority of those studies were performed using a verbal classification of the colors of the words placed in the screen (Siegrist, 1997), and not in the computerized version of the test that measures reaction time with millisecond precision, or by investigating the subgroups of results for each subject (ie, the level of congruence and incongruity of words and colors and their correct and incorrect answers). Yet, we found no evidence using scalp source activity using EEG in single-trial accuracy and SC recorded at the same time as the Stroop Test, nor measuring the cognitive effects observed in the test immediately before and after any type of meditation. This supports our evidence although it is necessary to increase the sample size of the study in future investigations. Another interesting point that can be observed, in addition to the randomization of the sequence of words and colors being unique for all the times the experiment was carried out, is that the results about the P300 are individual and seem to be linked to the time that each reported case practices the AST meditation. This does not corroborate the possible learning or retest effects of the experiment, since if there were, the learning effects would exhibit similar cortical activity due to a repeated performance of the same experiment (Gasser et al., 1985; McEvoy et al., 2000).

This indicates that the immediate effects of AST are linked to an enhancement in cognitive performance - measured through behavioral analysis of the Stroop test (Figures 3A, 5A and 7A) — with unique signatures observed in the amplitude and latency of the parietal and frontal P300 and also of the SC. This suggests a different central and autonomic immediate plasticity mechanism after AST meditation for each case that leads to a shorter response time in the test. Future studies with a larger sample are needed to confirm whether these results are linked to meditation time or individual differences, or to daily variations due to the current situation of the meditator, such as a stressful or relaxed day, or health conditions.

### 4.2. Study 2 - Characterization of frontal EEG synchrony during AST in traditional and wireless EEG

In this study we investigate the validation of a portable wireless electroencephalography (T-EEG) device during AST meditation. To verify whether the previous findings on alpha1 (8-10 Hz) frontal interhemispheric connectivity during AST meditation (Dillbeck & Bronson, 1981; Orme-Johnson & Haynes 1981; Travis et al., 2010; Tomljenović et al., 2016;) are found in the W-EEG, we performed a researcher-blind and randomized study for volunteers who performed three tasks on both T- and W-EEG’s for 20 minutes with eyes closed each during resting state (RS), cognitive activity (CA) — counting backwards from 1000 to 0 in steps of 4 —, and AST meditation. In addition to alpha1 band connectivity, we also investigated beta-band frontal interhemispheric coherence (13-30 Hz), which plays an important role in performing a cognitive task that requires mathematical calculations (Micheloyannis et al., 2005; Pierou & Vlamos, 216), as used in the CA condition of this study. Thus, we hypothesized that both devices would show the same properties in alpha1 and beta for the three experimental conditions performed by the subjects, with higher coherences in alpha1 for the AST condition and than beta in CA. The results confirm our hypothesis only for the alpha1 band, since a peak in beta frontal coherence was not observed during the performance of CA, but in the AST condition for both devices. For all reported case studies, the results are similar when comparing the three experimental conditions between the T- and W-EEG, ie, a trend towards greater frontal coherence in alpha1 and, surprisingly, higher beta during AST meditation. Our findings are important because they show that the W-EEG can reproduce the same properties found at the level of the same volunteer, indicating that future research can be carried out outside a controlled experimental environment to describe in more detail how this dynamic occurs during daily practice of the meditators. Previous studies have described the phase coherence across different experimental conditions with the Emotiv X, such as the interhemispheric dynamics in bipolar disorder (Handayani et al., 2017), emotional responses (Fattouh, 2016), mild cognitive impairment (Handayani et al., 2018). In the context of meditation, Jadhav et al., 2017 showed an emotional regulation after focused attention meditation using band-specific asymmetry, while Phutke et al., (2019) investigated the brain connectivity pre and post 8 week mindfulness meditation protocol and found evidence of increase in parietal interhemispheric connectivity after the protocol. However, to date, no studies using W-EEG were found on immediate effects of AST or the validation across different mental states with eyes closed as we investigated in this study.

Differentiating at the single-subject level different mental states performed with eyes closed is a challenge in the neuroscience field, as brain patterns with eyes closed when performing the same task are very subtle and highly influenced by alpha and mu synchronization in the occipital and motor cortex, respectively (Babiloni et al., 2010; Barry & Blasio, 2017). Bashivan et al., (2016) provides the only evidence in the context of the identification of different mental states with the W-EEG, however, they described the inherent dynamics with tasks being performed with eyes open. Thus, despite the size of our sample and the nature of the study, the results observed in the interhemispheric coherence of the alpha1 and beta bands during three mental conditions with closed eyes indicate that 1) the W-EEG can be a robust system for identifying states. of AST meditation, and 2) the alpha1 and beta coherence between the F3 and F4 electrodes can be an electrophysiological marker in identifying different mental states when the eyes are closed, especially during AST meditation.

An important point about the W-EEG is the coherence scale of all conditions in both frequency bands. Despite the trend of replicability of the results observed in the T-EEG, a decrease in synchronicity seen by the smaller scale of coherence for W-EEG (Figures 4, 6 and 8), which may be an indicator of 1) differences inherent in sending analog signals digitally via bluetooth, (2) wireless EEG system uses sequential data sampling (3) sampling rate, 4) not applying common averaging reference (CAR) - for validation purposes, we chose to use the same analysis pipeline for T- and W-EEG, however it is suggested that different pre-processing steps can be performed on the wireless device to find the same result at scale -, and 5) inherent hardware properties such as battery and signal amplification circuitry that are slightly different from T-EEG.

**Figure 4.**
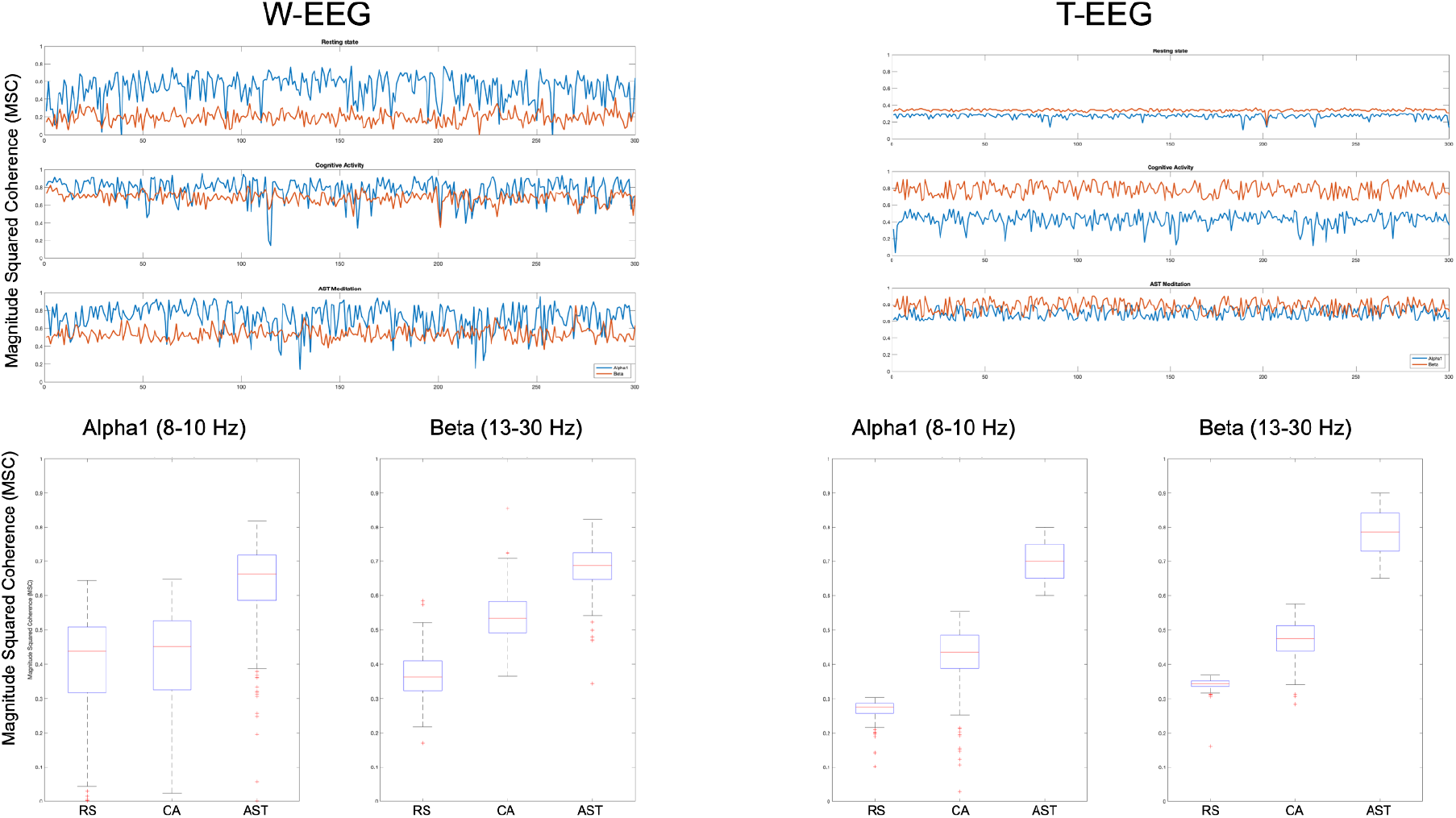
Frontal (F3-F4) phase coherence results for alpha1 (8-10 Hz) and beta (13-30 Hz) during the three experimental conditions performed randomically in traditional and wireless EEG’s for the case 1.

**Figure 5.**
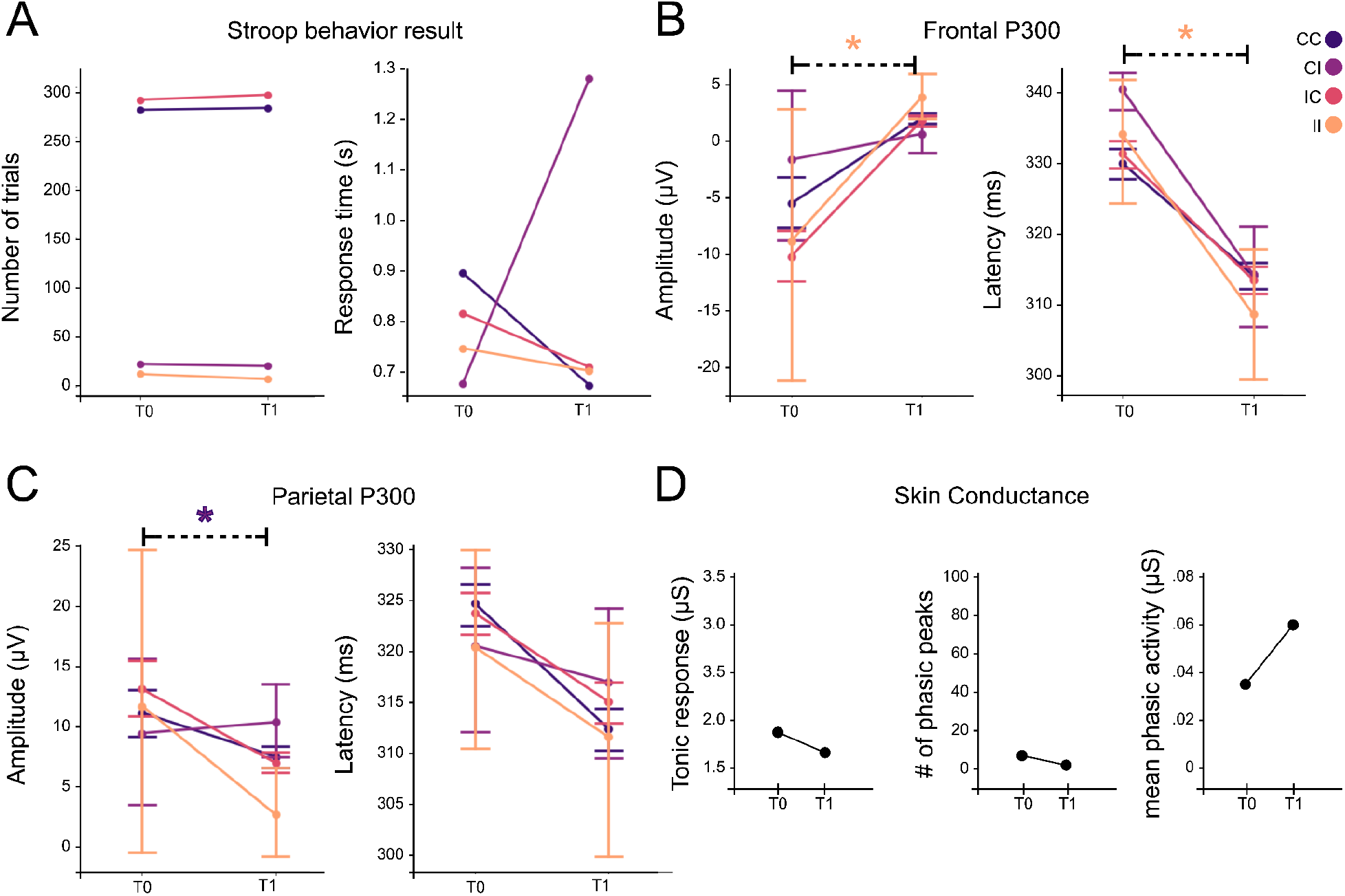
Case 2 results for experiment 1. The colors represent the Stroop task conditions (CC - Congruent Correct, CI - Congruent Incorrect, IC - Incongruent Correct, and II - Incongruent Incorrect). * show the statistical significance (two-tailed student’s T test corrected for multiple comparisons) < 0.05. The error bars represent the standard error of the mean for each condition.

**Figure 6.**
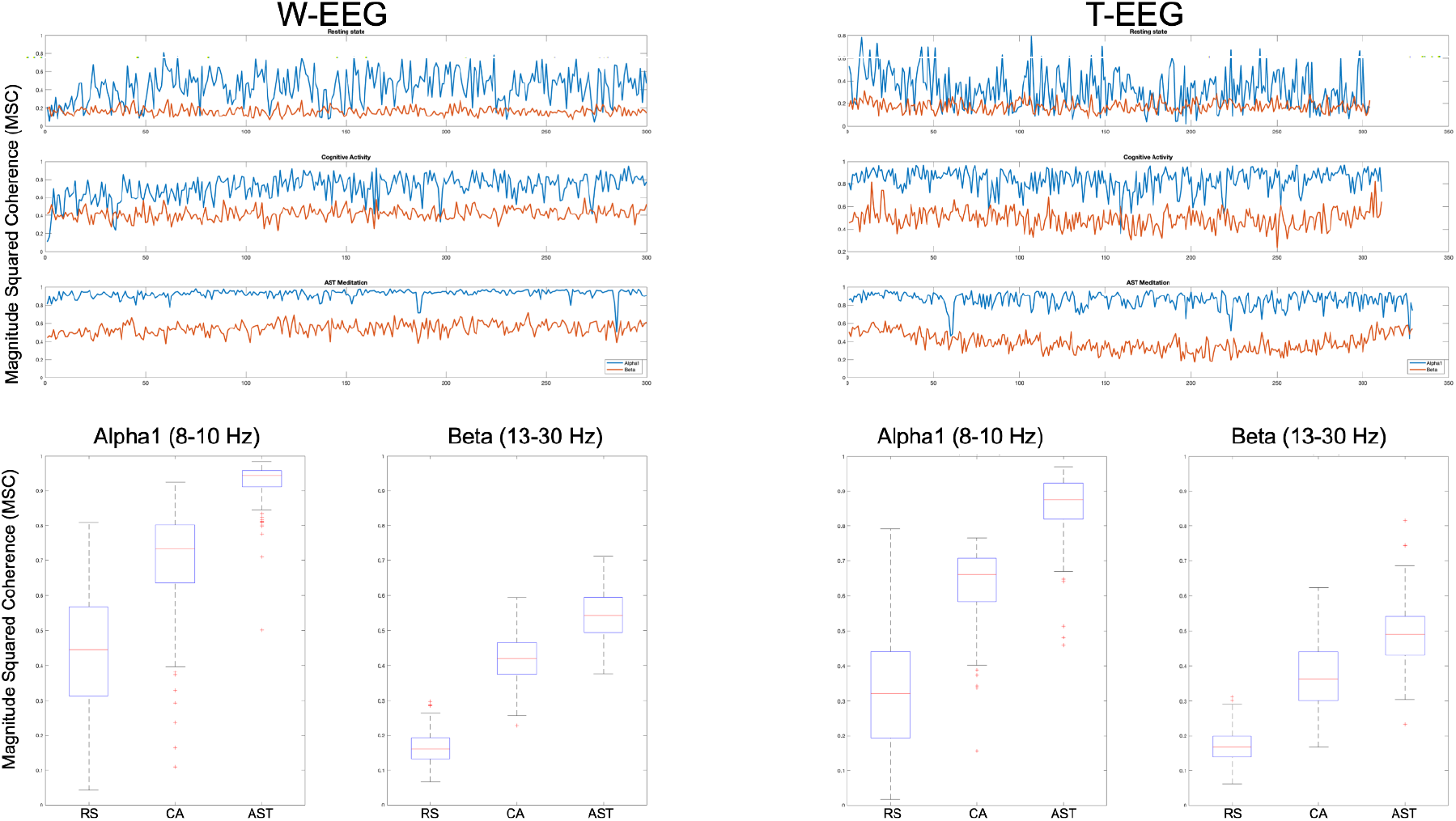
Frontal (F3-F4) phase coherence results for alpha (8-10 Hz) and beta (13-30 Hz) during the three experimental conditions performed randomically in traditional and wireless EEG’s for the case 2.

**Figure 7.**
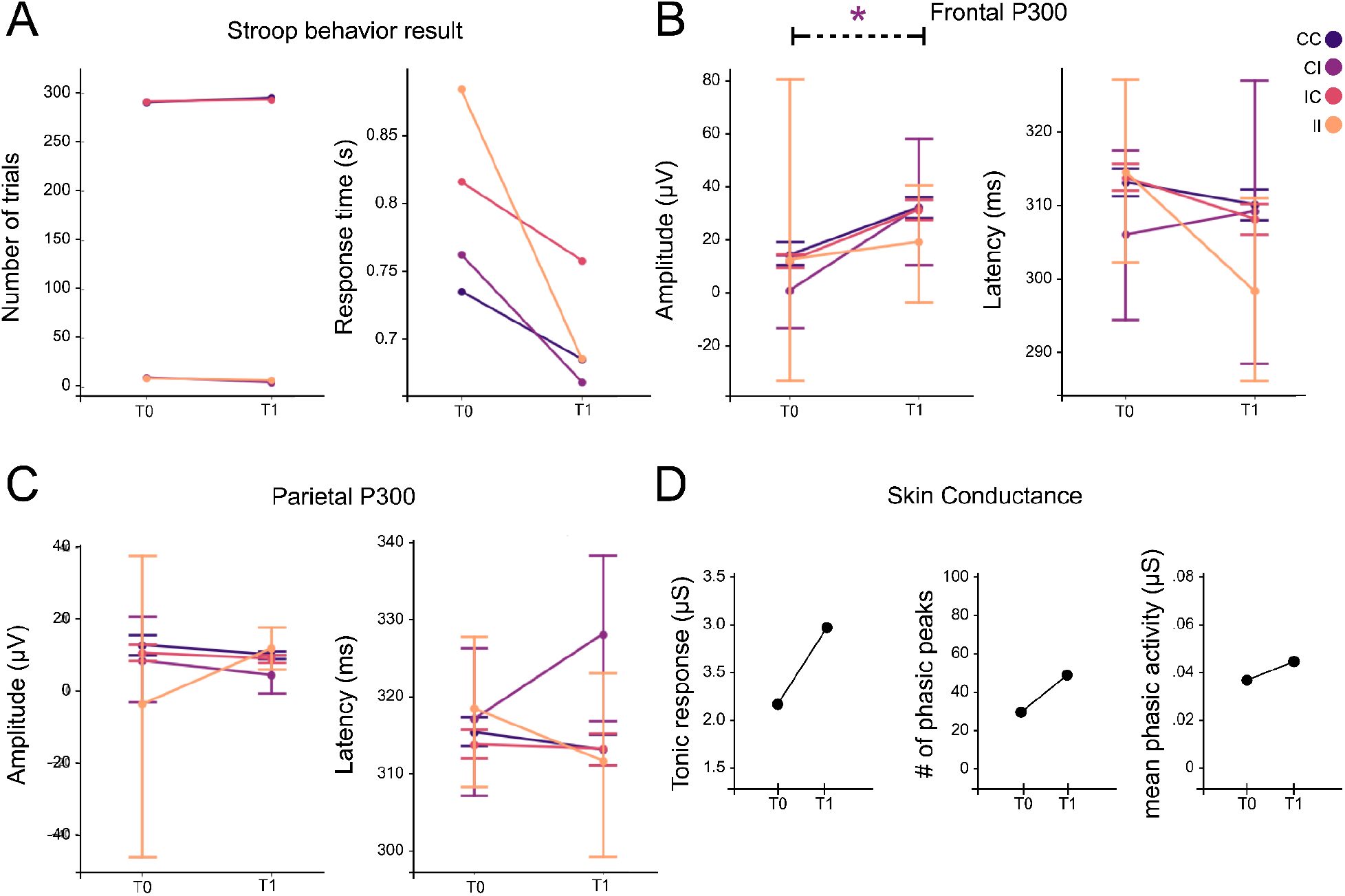
Case 3 results for experiment 1. The colors represent the Stroop task conditions (CC - Congruent Correct, CI - Congruent Incorrect, IC - Incongruent Correct, and II - Incongruent Incorrect). * show the statistical significance (two-tailed student’s T test corrected for multiple comparisons) < 0.05. The error bars represent the standard error of the mean for each condition.

**Figure 8.**
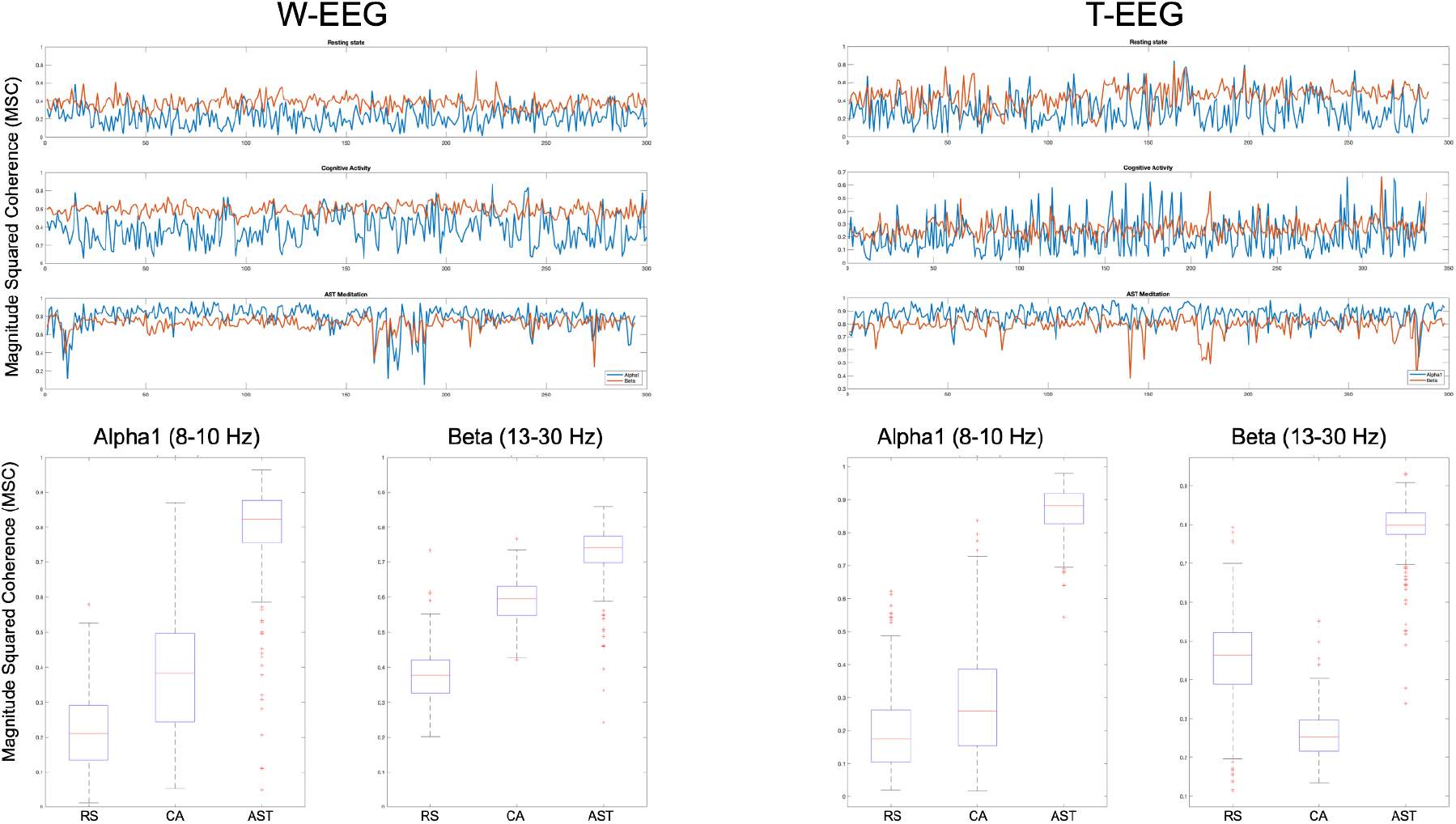
Frontal (F3-F4) phase coherence results for alpha (8-10 Hz) and beta (13-30 Hz) during the three experimental conditions performed randomically in traditional and wireless EEG’s for the case 3.

## 5. Conclusions

As discussed above, the main findings of this pilot study show that there is an immediate effect after AST meditation at the level of the same individual with different patterns of P300 and SC activity and that AST meditation is marked by an overall increase in the frontal coherence of alpha1 and beta bands, when compared to other mental states. In this context, Alonso et al., (2015) described that a higher beta coherence is found under the Stroop task, and Basharpoor et al., (2021) showed that the higher alpha and beta interhemispheric coherences are correlated with the behavioral performance in an executive function task. Thus, the results observed in study 2 show evidence that may explain the behavioral improvement observed during the Stroop Test (Study 1), since our results show that AST meditation is marked by an overall increase in alpha1 and beta band synchrony in the frontal region. We postulate that in addition to AST meditation being able to lead to an altered state of consciousness that has a non-dual characteristic, it is also able to modulate the attention, executive control and cognitive systems in an immediate way independent of the length of time practicing AST and exposure to the practice. We conclude that 1) there is an immediate effect on cognition and executive control after AST meditation, 2) the frontal interhemispheric coherence of alpha1 and beta bands are increased during AST, and 3) W-EEG exhibits the same characteristics observed in T-EEG and therefore can be used to describe cortical dynamics during AST. This is a pilot study that indicates early evidence of individual-level W-EEG stability to describe the effects of AST meditation. Future studies with a larger sample in a natural environment using W-EEG are needed to confirm this hypothesis.

## Acknowledgements

We thank all the subjects who participated in the study. This work was supported by Coordenação de Aperfeiçoamento de Pessoal de Nível Superior (CAPES), and Conselho Nacional de Desenvolvimento Científico e Tecnológico (CNPq).

## Author contributions statement

L.G. designed, conceived and conducted the experiment, analyzed the data and wrote the manuscript, G.M.S. and T.A.S.B. conducted the experiments and revised the manuscript, N.A.S revised the manuscript and provided the experimental setup, D.O.J interpreted results, wrote and reviewed the manuscript. All authors contributed to the manuscript.

## Data availability statement

The data that support the findings of this study are available on request from the corresponding author, Lucas Galdino.

## Supplementary Material

**S1.**
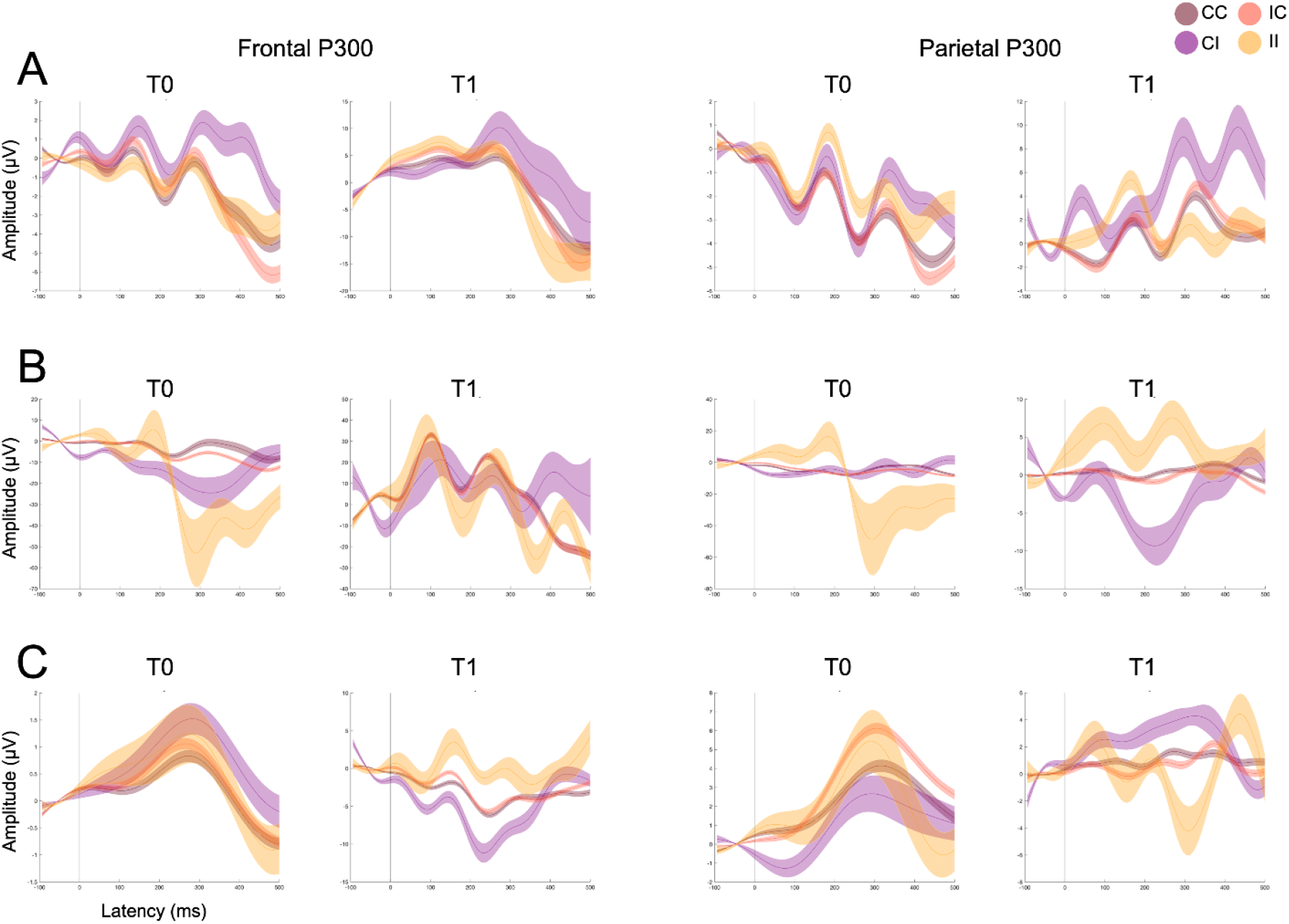
Event-related Potentials for the case 1, 2, and 3 shown in A, B and C, respectively.

